# Improving single molecule localisation microscopy reconstruction by extending the temporal context

**DOI:** 10.1101/2025.04.05.647262

**Authors:** Sebastian Reinhard, Vincent Ebert, Jann Schrama, Markus Sauer, Philip Kollmannsberger

**Affiliations:** Department of Biotechnology and Biophysics, University of Wuerzburg, Am Hubland, Wuerzburg, 97074, Bavaria, Germany; Biomedizinische Physik, Heinrich-Heine-Universität Düsseldorf, Universitätsstr. 1, Düsseldorf, 40221, Nordrhein-Westfalen, Germany

**Author notes:** Contributing authors.

## Abstract

Single-molecule localization microscopy methods such as *d*STORM require specific buffer conditions to enable blinking and detection of individual emitters, making them incompatible with live cell imaging and expansion microscopy. An alternative approach to achieve super-resolution without blinking is to observe the fluctuations of the emitter intensity over time. Existing localization algorithms for high-emitter density make use of radial symmetry or use artificial neural networks trained on single high-density frames to predict emitter positions. Here, we aim to improve the resolution by using a larger temporal context. We combine the U-Net architecture used previously for image reconstruction with multi-head attention used in the Transformer architecture. We compare the results to DECODE and eSRRF as well as to traditional fitting algorithms on public benchmark data. A generic pre-trained model is provided together with a fast and robust simulator for training data and all scripts needed to train custom networks.

## 1 Introduction

### 1.1 Background

Single-molecule localization microscopy (SMLM) based on *direct* Stochastic Optical Reconstruction Microscopy (*d*STORM) [1] is a widespread method to image nanoscale macromolecular structures in cells and tissues. Its resolution is superior to other super-resolution light microscopy techniques such as STED [2] and SIM [3], but is ultimately limited by labeling density, linker size and crosstalk between nearby fluorophores [4]. The underlying principle is to stretch the signal over time by introducing conditions such that emitters are only active during a fraction of time, and to then reconstruct the positions of individual emitters in diffraction-limited frames. The most widely used reconstruction algorithms are based on fitting radially symmetric Gaussian distributions to diffraction-limited spots in single frames to predict the coordinates of individual localizations. Many open source software packages are available that focus on different aspects such as speed, precision, or ease of use [5–7]. Under ideal conditions, this approach has been shown to approach the theoretical Cramér-Rao lower bound (CRLB) for inferring the parameters of a distribution from noisy data.

The state of the art of SMLM fitting is monitored in the SMLM fight club, a benchmark with a variety of datasets for different imaging conditions, sample types and densities, based on standardized evaluation measures [8]. Recently, localization algorithms using artificial neural networks (ANNs) have surpassed all other methods in terms of versatility and precision [9, 10]. ANN-based fitters either learn a mapping from the raw frames to super-resolved images (DeepSTORM), or predict the parameters of a Gaussian distribution (position and uncertainty) for each localization (DECODE). The latter method outperforms all other approaches when trained on simulated data with the same imaging parameters and noise distribution as the data to be predicted. ANN-based methods can also be trained to learn unconventional point spread functions (PSFs) [11, 12].

Improving the reconstruction of SMLM data is an ongoing field of research. Typical limitations of current approaches are that they require well-separated localizations, sufficient signal-to-noise ratio, and only look at individual frames by default. Not only is the long acquisition time a limitation, but blinking requires specialized imaging buffers that are incompatible with live imaging and expansion microscopy [4]. To apply localization-based super-resolution microscopy in these scenarios, robust reconstruction algorithms for high emitter densities with less blinking are needed.

### 1.2 Temporal context

While the above mentioned methods process individual frames or up to three consecutive frames, an alternative strategy is to use correlated temporal fluctuations in the signal as source of information. This concept was introduced with SOFI by Dertinger et al. [13]. The intensity of individual pixels is correlated over time due to the blinking of the emitters. The higher-order autocorrelation functions contain higher powers of the PSF, resulting in improved resolution. Cross terms in the correlation that would lead to artifacts are eliminated by calculating cumulants (integrals over correlation functions). The auto-cumulants of independently treated pixels can quickly be calculated, whereas the cross-cumulant takes spatial correlations between pixels and overlapping PSFs of nearby emitters into account to generate additional “virtual” pixels, but is more complex. The static background in the resulting images is highly suppressed, making these images well suited for subsequent deconvolution compared to noisy images. Limitations of this method are the increasing computational complexity with increasing cumulant order, and the sensitivity to fluctuating noise. Furthermore, the molecular brightness of the dyes also appears in higher orders in the cumulants, resulting in the suppression of dim emitters. In practice, SOFI is limited to fourth order cumulants, leading to a two-fold improved resolution compared to widefield imaging.

SRRF [14] uses a radiality gradient filter to enhance local intensity peaks. Noise is then suppressed by calculating temporal cumulants as in SOFI, but on the radiality maps instead of the intensity images. This alleviates the dependence on intensity in the original SOFI method, where bright emitters can overshadow dim ones due to the nonlinear intensity dependence in the higher-order cumulants. The same approach is followed in eSRRF [15], but with better interpolation and parameter selection as well as a detailed quantification of the performance.

A different class of methods that shows good performance at high densities is compressed sensing (CS) [16]. Rather than predicting individual emitter coordinates, the image is projected onto a high-resolution pixel grid by inverting the imaging process. In real scenarios under noise and without exact knowledge of the PSF, this approach requires regularization. In CS, regularization is based on assuming sparsity of the underlying image, i.e., only a very small fraction of pixels in the reconstructed super-resolved image contains non-zero intensities.

CS can also use temporal context by assuming sparsity in the fluctuation domain [17]. SMLM measurements are not only sparse in space with only a few emitters active per frame, but also in time, since emitters are in an non-fluorescent off state most of the time. Each pixel in the high-resolution space is only active within a fraction of the total time. The implementation in SPARCOM using the FISTA iterative algorithm shows good results, but is prone to the same limitations as other CS-based methods, such as large computational demands, thinning effects due to the imposed sparsity, and parameter sensitivity. A promising development is the unrolling of the iterative optimization behind compressed sensing into a series of convolutional neural networks ([11, 18]).

#### 1.2.1 Our approach

Since the publication of SOFI, the field of fluctuation-based super-resolution has undergone much development, as summarized in [19]. While there is still a lot of potential behind fluctuation imaging, existing methods do not make full use of its possibilities. SOFI is confined to lower orders in practice, while localization fitting with rapidSTORM or DECODE is limited to 2-3 consecutive frames. CS makes assumptions about sparsity, which also limits its practical use. In principle, the intensity of a single emitter is correlated over long time spans and many frames. A trainable algorithm could learn to extract both spatial and temporal information when presented with a noisy signal. The key requirement to successful deep learning fitting using temporal context would be a good simulator for training ground truth with detailed information about the noise and the scales of the signal.

Here, we present a new SMLM fitting method that combines convolutional neural networks with multi-head attention as used in the Transformer architecture [20] for extended temporal context. We developed an accurate simulator for EMCCD-based SMLM experiments to generate ground truth. The performance of our approach at different localization densities is demonstrated on simulated data with known emitter positions, as well as on public benchmark data. We compare the results to the CRLB for our method as well as for several state-of-the-art tools. Ablation studies are conducted in order to investigate the importance of different parts of the architecture. Where possible, we adopted well-established techniques, such as the spatial encoding of emitter positions and the GMM-based loss from DECODE.

All of the previously described methods (with the exception of SPARCOM) do not use an extended temporal context and only process 1-4 frames at a time. Every emitter is typically active multiple times during an experiment or over a larger number of frames in the case of high density imaging. Given that the emitter position does not change over time while the background intensity and activity of other emitters do, there is relevant correlated information in the time context that can be used to infer the emitter position. This information is difficult to formalize but could be extracted by a neural network if fed with sufficient time context.

Neural network architectures that can process temporal context are used in the field of natural language processing [20, 21]. Current state-of-the-art approaches are based on the attention mechanism, which, although it has been adapted to process image embeddings [22], is naturally suited for sequence data. A combination of neural-network-based image reconstruction with attention-based temporal context would be an obvious approach to improve localization microscopy. A drawback of Transformers is that they require extremely large training datasets and a large number of parameters, rendering the training process prohibitively slow and expensive in cases where re-training for specific imaging scenarios is needed.

Here, we explore the combination of U-Net fitting with an intermediate attention block that processes temporal context. Since PSFs are typically small, we use a shallow spatial compression, but a large temporal context. This leads to a slim architecture that allows retraining with low resource requirements.

## 2 Results

### 2.1 Simulations

In order to train and validate an artificial neural network to correctly predict the position of the emitters in SMLM, a ground truth with known emitter positions that accurately mirrors real experimental data is required. We thus started by developing a simulation engine that generates ground truth raw frames at high rates with accurate photophysics and camera noise and with tunable parameters to adapt to changing conditions. Here we place particular emphasis on the changes over time, which are crucial for temporal correlations.

The blinking behavior of an emitter is defined by multiple state transitions and their lifetimes. A singlet-singlet transition can lead to the emission of a photon, while other processes lead to non-fluorescent time intervals. In classical SMLM, the aim is to keep the emitter in an ON state for the duration of one frame, followed by an OFF state for a longer time interval. This leads to the desired sparse distribution of emitters. However, these conditions are not always experimentally feasible. Longer ON times lead to higher emitter densities, complicating the reconstruction. The encountered longer ON times can, on the other hand, be a chance to yield an increased reconstruction precision with fewer frames since more photons are emitted over multiple frames. A realistic switching behavior can be modeled with a Markov Chain as explained in [23]. A detailed description of our adaption is shown in Fig. 8 and described in the methods section.

Another important component of microscopy images is the camera characteristics. Due to their high sensitivity, EMCCD chips make up a significant fraction of the cameras used in SMLM. Therefore, we focus our model on this type of camera. The underlying noise characteristics are a joint model of Poisson, Gamma and Normal distributions and are well described in literature [24–26]. Poisson noise occurs during discretization when photons are converted to electrons on the camera chip. Gamma noise originates in the photon multiplication register, and normally distributed readout noise is caused by the conversion of electrons into a digital signal.

The third key component is the background signal caused by autofluorescence or unspecific labeling. This background can lead to conditions that overstrain algorithms based on local maxima detection (MLE, fitting), thresholding (CS), or gradient detection (SRRF). We create this background by adding random patterns consisting of random integers in the range of ∈ [0, 255] convolved with a Gaussian kernel of 5 *px* width and height.

This simulation engine forms a necessary part of the reconstruction workflow, enabling the adaptation of the network to various experimental scenarios and measurement conditions.

### 2.2 Neural Network with temporal context

Our goal is to extract both spatial as well as temporal information to predict emitter positions. The architecture of an artificial neural network should be designed to constrain the search space and to be able to take a series of input frames and predict emitter positions. The fitting of emitters in one frame is a redundant and locally constrained problem. Convolutional neural networks, applying the same operation to patches all over the image, are therefore well suited to collect the spatial information necessary for a precise fit. This concept is already applied in current deep learning fitters based on the U-Net architecture [9, 10]. The temporal component of emitters switching is however much more complex and only sparsely accessed in current architectures. Switching occurs over long periods of time and requires a large context size to access information of multiple blinking events. We combined the U-Net architecture with the Attention mechanism to encode sequential information, as used in the Transformer architecture.

Our architecture consists of three parts: an initial U-Net [27] embeds spatial information from individual frames and outputs a feature map with the same size as the input images. We then use the processed spatial information as input tokens for our attention module. The output of the attention module is an encoding of the temporal context in the input sequence. The second U-Net maps this information onto a multichannel output image containing the probability of finding an emitter in the respective pixel, as well as its position, uncertainty, intensity and background.

The PSF of individual emitters in SMLM images usually spans only 5-8 pixels before the signal falls below noise levels. This means that the pixels outside an 8×8 pixel patch contain no relevant information about an emitter at its center. Assuming that the PSF is the same over the entire input image, the position of a patch within the input image contains no relevant information. We therefore use a shallow U-Net as an encoder for our network.

While the information in the spatial domain is highly constrained, the time context poses a more complex problem. Switching events occur over longer periods, including OFF frames with little to no information. To analyze the temporal domain, we implemented an attention block (Fig. 2b, see methods for details) [20] to collect information on a large number of frames. The attention mechanism takes a batch size of up to 50 frames and an embedded dimension equal to the hidden dimension of the U-Net. It is placed behind the first U-Net. The second U-net in our architecture aims to collect this temporal information. Similar to the first block, we compress spatial information in two downward convolution steps. The subsequent upward pass uses activated concatenation as described in [28]. The output contains twice as many feature maps as the input while the width and height remain unchanged.

**Fig. 1.**
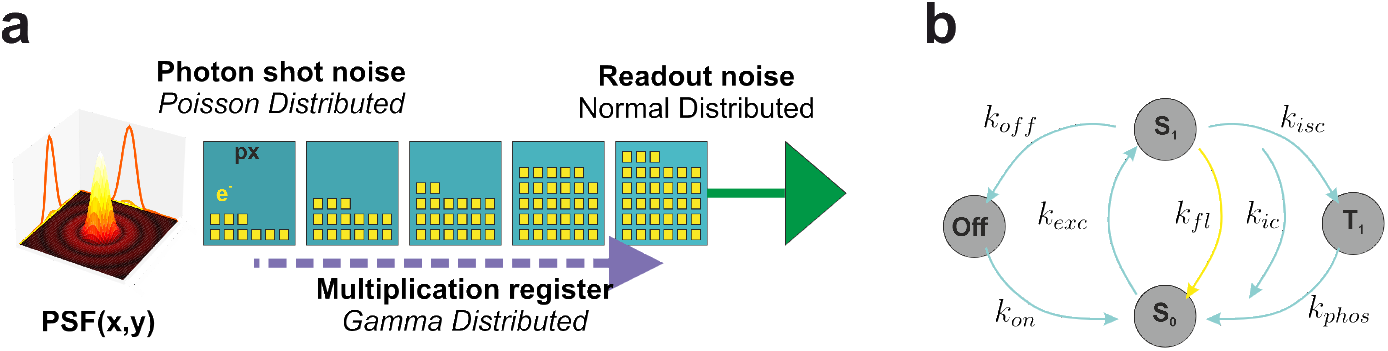
**a** Spatial simulation pipeline: The core of our simulation is the PSF. We sample the Poisson distribution of the expected number of photons (based on **b**) multiplied by the integrated probability over one pixel. Electrons are further multiplied in the multiplication register. The corresponding noise is Gamma distributed. Converting the analog into a digital signal adds normally distributed readout noise. **b** Simulation of photon traces is based on a Markov chain model with transition rates between the different fluorescent and non-fluorescent states.

**Fig. 2.**
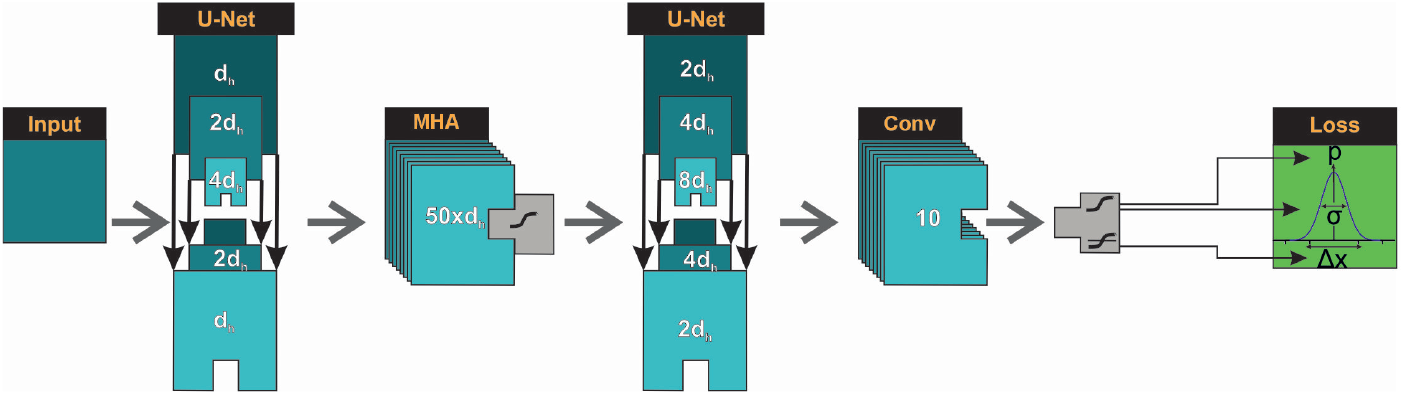
Network Architecture. The images are initially processed with an U-Net, increasing the feature size to the number of hidden dimensions *d*_*h*_. Subsequently we apply a positional encoding and a multihead attention block over batch and feature dimension. A second U-Net processes the additional temporal information and produces 2*d*_*h*_ feature maps. We apply 4 different convolutional heads to extract information for probability, mean values of Gaussian distributions, standard deviations of Gaussian distributions and background information into 10 final feature maps.

**Fig. 3.**
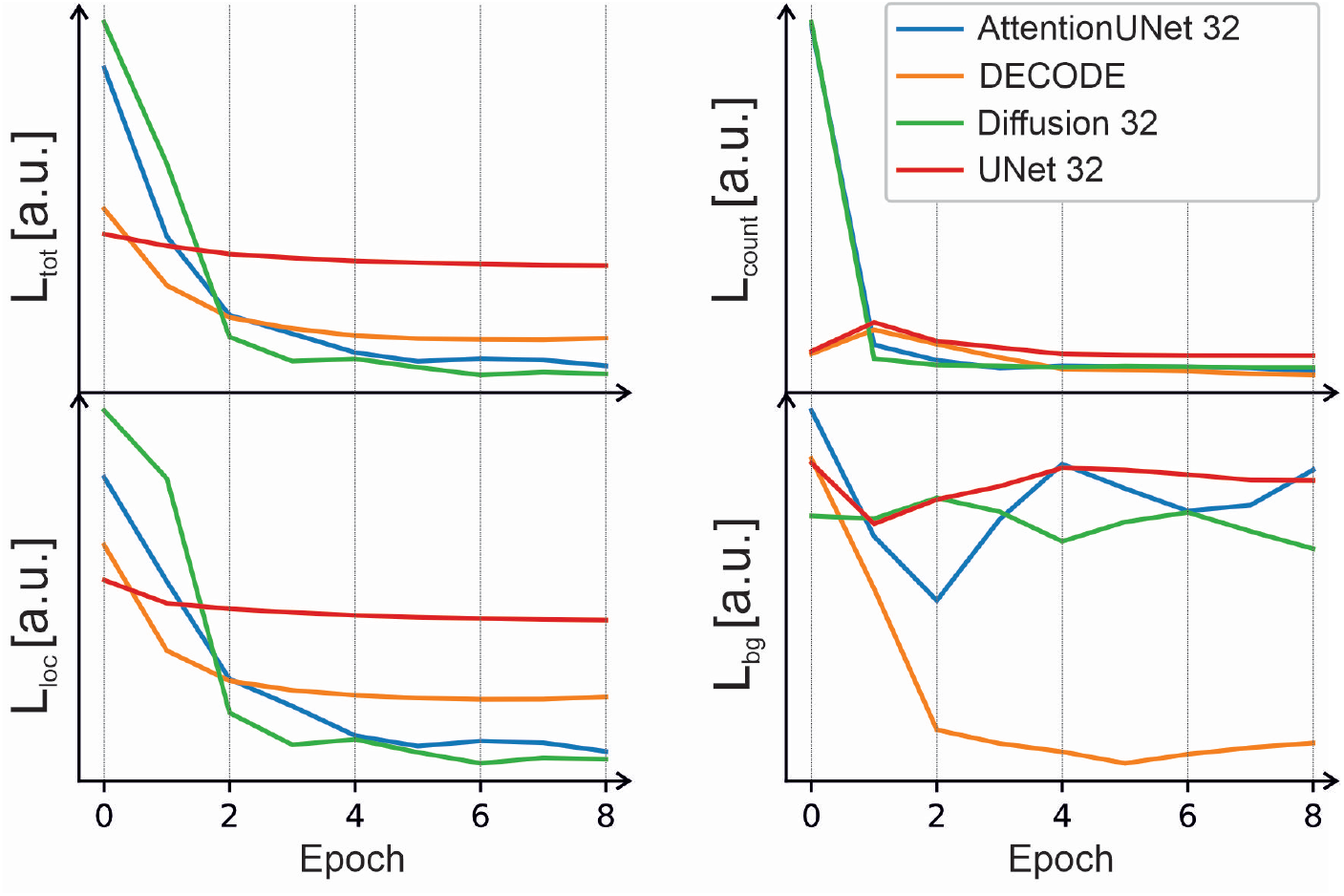
Validation loss components of the evaluated networks over 20 training iterations and 9 evaluation epochs. Top left: total loss, top right: count loss (eq. 20), bottom left: localization loss (eq. 18), bottom right: background loss (eq. 21). The hidden feature dimension was 32 in all cases.

We implement a convolutional head (3×3 + 1×1 convolution) for each activation function to construct a suitable number of output feature maps, followed by a final activation that constrains outputs to reasonable intervals for the loss model. Probability is constrained to ∈ [0, 1] with a sigmoid function, mean values are limited to ∈ [−1, 1] with a tanh activation, and standard deviations are restricted to ∈ [0, 3] with a sigmoid activation multiplied by 3. The background map remains unconstrained.

### 2.3 Hyperparameter selection and ablation tests

The choice of hyperparameters and the different parts of the network may have an influence on performance. To better understand the importance of the different components for the results, we systematically varied the hyperparameters and performed ablation studies by removing different parts of the network. All networks were trained for 20 iterations on the “Base” dataset (Table 7) and evaluated with a separate validation dataset with similar parameters. The performance is recorded every two iterations. We evaluated the architectures listed in Table 1. The Diffusion network is described in 5.4. Adding a Multilayer Perceptron after the attention block did not significantly improve the performance. The second U-Net on the other hand increases the positional performance significantly. Positional encoding of the input frames had no positive effect on the validation loss, and an attention mechanism without a second U-Net only improves the background loss. Increasing the depth of the U-Net leads to overfitting. When directly embedding the input patches without the first U-Net, the positional information gets slightly worse.

**Table 1.** Comparison of the different tested network architectures in terms of parameters and performance.

|  | UNet_32 | DECODE | AttentionUNet_32 | Diffusion_32 | AttentionUNet |
| --- | --- | --- | --- | --- | --- |
| Hidden dimension | 32 | 48 | 32 | 32 | 98 |
| Context length | 1 | 3 | 50 | 50 | 50 |
| Efficiency (eq. 23) | 49.1 | 71.4 | 78.1 | 77.8 | 79.6 |
| Number of parameters | 558262 | 2199418 | 2797750 | 4329270 | 25117110 |

We observed that fine-tuning a trained model works much better than training each condition from scratch. Training a new network on dataset “CRLB 4” achieved an efficiency (Eq. 23) of 52 after 600 iterations, whereas fine-tuning on the pretrained network achieved an efficiency of 91 after only 60 iterations.

### 2.4 Proof of concept

When fitting a distribution to measured data, the precision of the fit is fundamentally limited by the information contained in the measurements. This limitation is quantified by the Fisher information, which represents how much the data informs a particular fit variable. Fisher information is additive, indicating that it scales linearly with the amount of available data points. It is directly related to the Cramér-Rao Lower Bound (CRLB), which provides a theoretical minimum for the uncertainty of an estimated parameter.

In the context of localization microscopy, the number of detected photons *N* defines the amount of available information. The CRLB thus sets the lower bound on the uncertainty of estimating the distribution’s mean, *µ*, by the photon count. For a noiseless

Gaussian fit of a continuous space, the uncertainty of *µ* is given by:

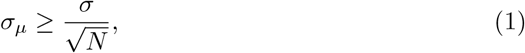

For our estimations, we use a more sophisticated term taking the discretization of the grid into account:

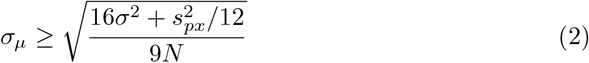

where *s*_*px*_ denotes this pixel size (taken from [29] with background intensity *b* = 0). While this term takes a discretisation of the Gaussian function through pixels into account, it does not compensate for noise sources of the camera simulation.

The CRLB can be used to test the benefit of the extended temporal context: if information from different ON states is combined, the statistic should be improved to a point where the fits are far below the CRLB of one frame. To test this, we simulated a dataset with multiple switching events over the 50-frame context and performed a low-density fine-tuning for DECODE and Attention U-Net (Dataset 7: “CRLB 4” for training and “CRLB 3” for testing). We used a train-test split of 80-20 and optimized the hyperparameters with optuna [30] for 20 trials, which resulted in an optimal efficiency of 91 for DECODE and 94 for AttentionUNet. In Figure 4, we plotted the CRLB for emitters in a single frame, the CRLB of all combined ON events, and the distance of the fit to the ground truth. While the 3-frame time window of DECODE already outperforms the CRLB of one frame used by ThunderSTORM, the AttentionUNet yields even lower distances with 76.59 percent of emitters below the CRLB in comparison to 39.95 for DECODE and 17.53 for ThunderSTORM. Combining all fits for each emitter over the 50 frame timeframe, we collected additional metrics to gain deeper insights about what is happening under the hood of the attention mechanism. We computed the average fit distance and the corresponding standard deviation for each emitter. The averages of these metrics are shown in Table 2. We found the smallest dispersion for the AttentionUNet, which indicates that emitters are brought into context over the given timeframe.

**Table 2.** Evaluation of CRLB 3 dataset

|  | DECODE | AttentionUNet | ThunderSTORM |
| --- | --- | --- | --- |
| Best efficiency (eq. 23) on validation | 91 | 94 | — |
| % of undetected emitters | 0.18 | 1.99 | 4.89 |
| Averaged (mean distance) fit to emitter | 8.82 | 6.12 | 18.3 |
| Averaged (std distance) fit to emitter | 4.35 | 1.85 | 10.66 |
| % under CRLB of corresponding frame | 39.95 | 76.59 | 17.53 |
| JI overall | 0.96 | 0.95 | 0.77 |
| RMSE overall | 10.87 | 7.51 | 24.22 |

**Fig. 4.**
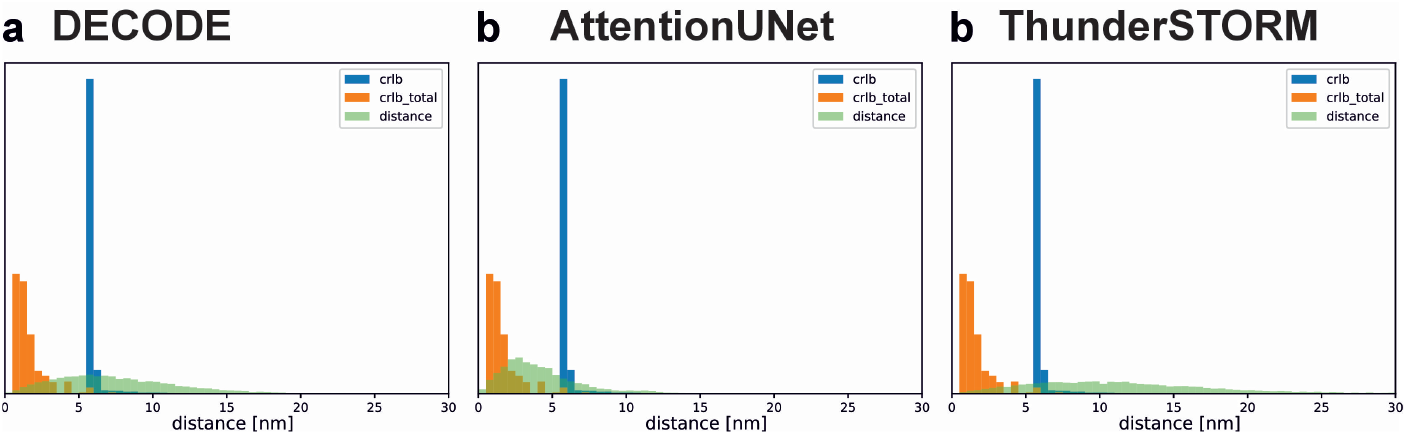
Histogram of the distance from fit to ground truth (green), the CRLB of each emitter per frame (blue) and the total CRLB of an emitter (orange) accumulated over the whole context.

### 2.5 High density reconstruction and generalisation

We next investigated how well the network performs under higher densities and how well a single training generalizes. For this, we used the model trained on the dataset “Base” (Table 7) with a density of 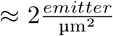. We evaluated using a dataset with localisations distributed over a binary image of our lab logo. This structure is well suited for evaluation purposes since it contains extended areas (in the lemon) as well as finer and more complex structures in the letters. We simulated densities of 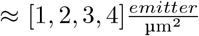 in the populated areas. Results are shown in Figure 5, and the corresponding metrics are listed in Table 3. While the AttentionUNet outperforms DECODE for “Density 2” and “Density 3”, DECODE achieves a better *JI* for “Density 4” and overall better metrics for “Density 1”. Since “Density 2” and “Density 3” are closer to the training data, one can assume that DECODE generalizes better. Taking the image of “Density 4” into account, a better *JI* and lower *RMSE* might indicate a higher scattering of fits for DECODE leading to a “fitting by accident” for highly populated areas.

**Table 3.** Evaluation of different emitter densities

|  | DECODE JI | DECODE RMSE | AttentionUNet JI | AttentionUNet RMSE |
| --- | --- | --- | --- | --- |
| Density 1 | **0.94** | **26.01** | 0.61 | 43.11 |
| Density 2 | 0.80 | 37.45 | **0.82** | **26.24** |
| Density 3 | 0.63 | 46.68 | **0.65** | **34.15** |
| Density 4 | **0.51** | 52.72 | 0.43 | **47.09** |

**Fig. 5.**
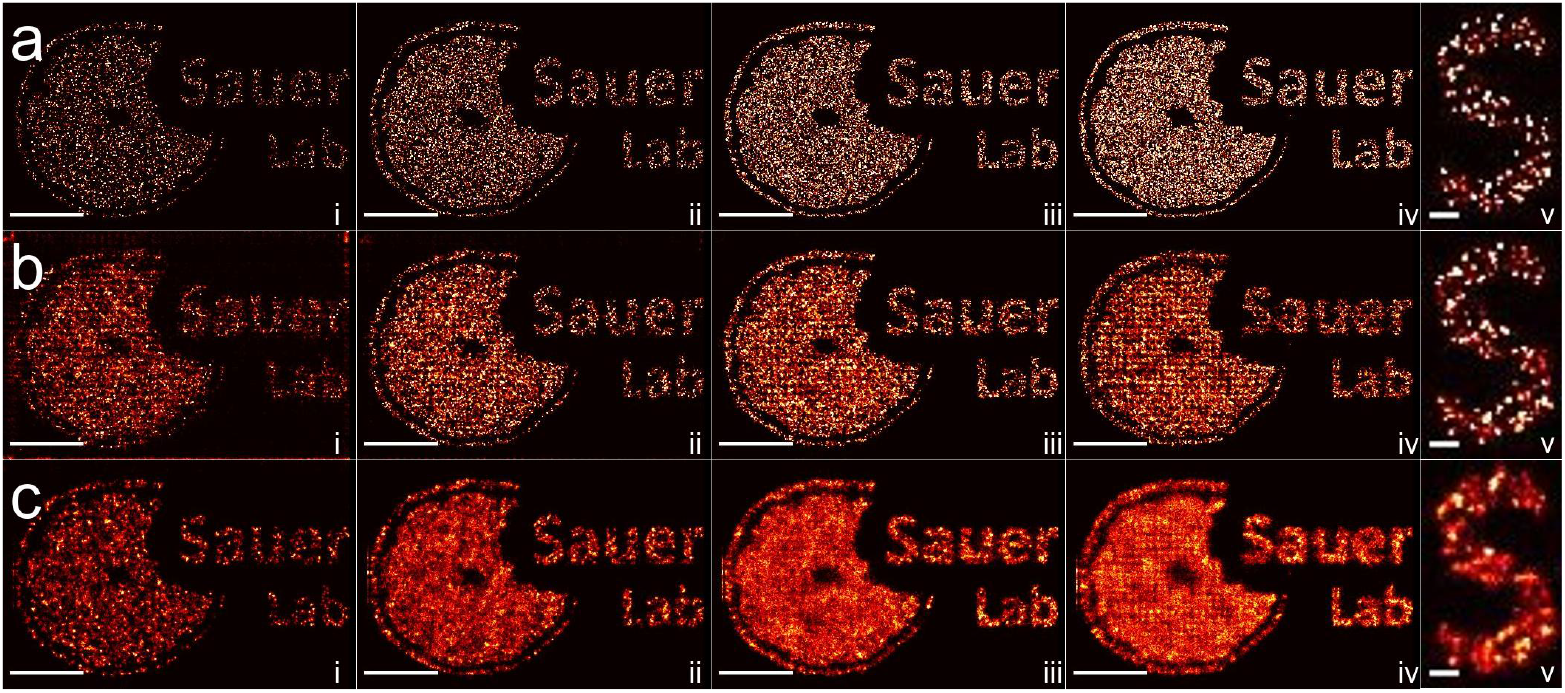
Network comparison under different structural densities from 1 to 4 emitters per µm^2^ (**i**-**iv**). **a** Ground truth. **b** AttentionUNet. **c** DECODE. Column **v** depicts a zoom in on density 2. An enhanced temporal context can provide additional information about the emitter position, resulting in an improved RMSE of our AttentionUNet compared to DECODE. Our network is however more susceptible to conditions that differ from the training density 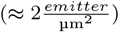. Scalebar: 100 nm for (**v**), 1 µm for (**i**-**iv**).

### 2.6 Visual performance and comparison with state-of-the-art algorithms

To perform a visual comparison of the reconstruction quality of our network with other state-of-the-art reconstruction algorithms, which are in parts not localisation based, we simulated a dataset with two approaching lines with a width of 60 nm and exponentially decreasing distance (dataset “Approaching lines”, Table 7). Figure 6 compares the AttentionUNet with SRRF, Compressed Sensing, DECODE and ThunderSTORM.

**Fig. 6.**
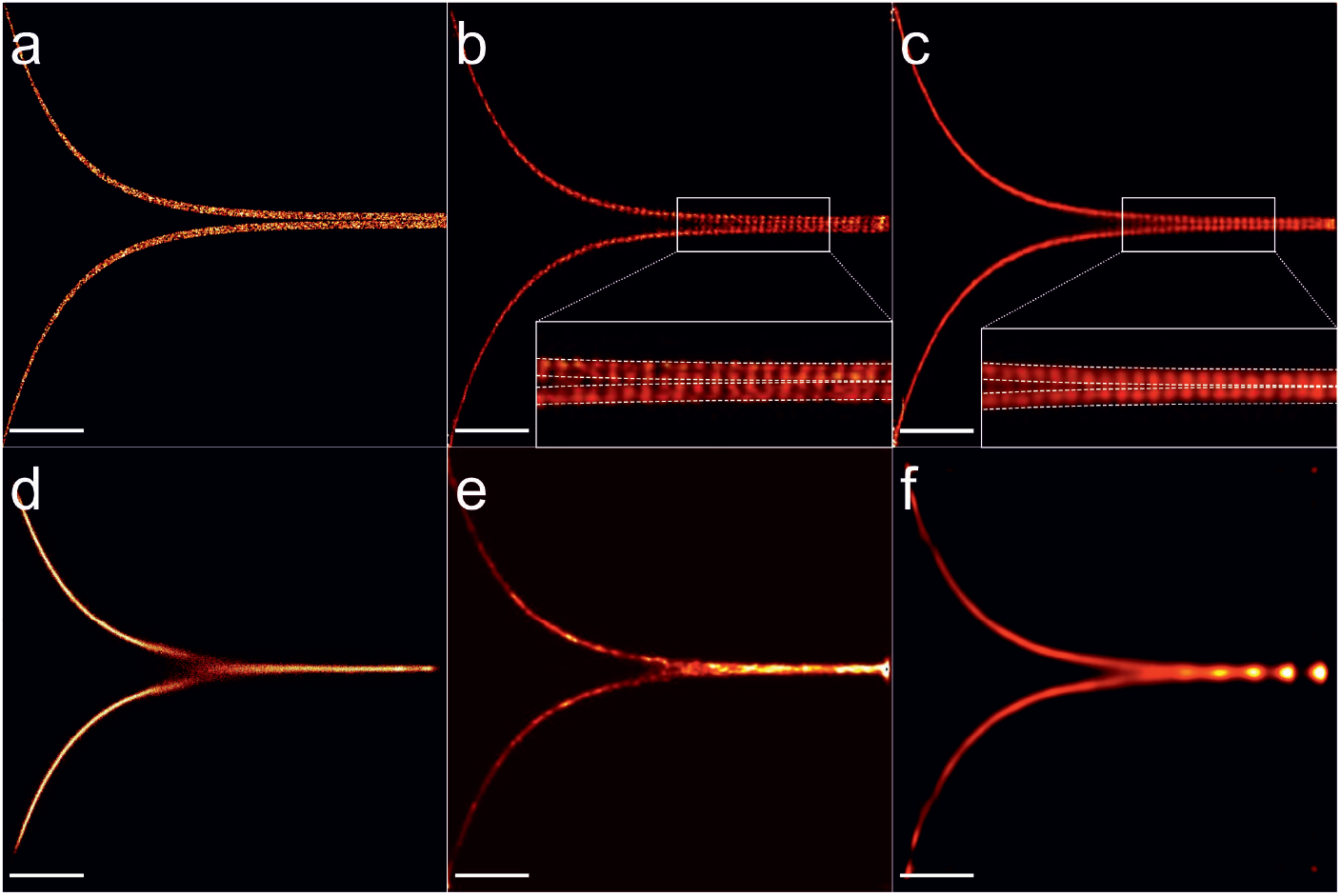
Simulated dataset of approaching lines. **a** Ground truth. **b** AttentionUNet. **c** DECODE. **d** ThunderSTORM. **e** SRRF **f** Compressed Sensing. The dotted lines in the zoom in (**b**,**c**) indicate the borders of the ground truth. Scale bars 1 µm

ThunderSTORM, developed for single emitter fitting, is unable to detect emitters in the region where the two lines approach each other. For the algorithm, overlapping emitters in this region appear as elongated in the y-direction and are therefore not detected. As the lines get closer together, multiple emitters are recognized as single ones with larger FWHM, leading to a line with reduced diameter.

Compressed Sensing pulls emitters apart by creating the sparsest possible solution under a given PSF and noise level. The result is in this case negatively impacted by the added background fluorescence, leading to artifacts at the right edge of the lines. SRRF applies an autocorrelation to the radial fluctuations of the image. The reconstruction is negatively impacted by noise. The lines merge much faster than in the ground truth and are depicted with a reduced width.

DECODE and the AttentionUNet, trained under similar conditions, provide the best reconstruction of the lines. While DECODE has a higher spread of the localisations (as we saw in the CRLB test), the lines look clearer in the AttentionUNet reconstruction. Both algorithms suffer from pixelation artefacts in the approaching regions. As shown in Table 4, both algorithms yield a similar *JI*. The AttentionUNet has however a much lower RMSE.

**Table 4.** Evaluation of Approaching lines Dataset

|  | DECODE | AttentionUNet | ThunderSTORM |
| --- | --- | --- | --- |
| JI | **0.47** | 0.46 | 0.13 |
| RMSE | 48.37 | **42.99** | 61.77 |

### 2.7 Super-resolution fight club

To provide a comparison for a dataset that is independent from our simulation tool, we used the high density dataset from the super-resolution fight club [8]. This dataset consists of 8 filamentous lines with a density of 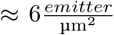. The PSF, with a sigma of *σ* = 109 nm in the focal plane, broadens up to a sigma of *σ* = 218 nm for 300 nm axial difference. Since we did not implement 3D PSF’s yet, we approximated the average FWHM to 150 nm and simulated a fine-tuning dataset for training (dataset “Contest(fine-tune)” in Table 7).

Reconstructions for AttentionUNet, DECODE, SRRF and CS are shown in Figure 7. SRRF is not able to reconstruct the overlapping areas in the high density regimes at the bottom left of the image. Additionally, artifacts are reconstructed between the individual lines. CS suffers from pixelation artifacts in regions where the PSF changes due to offsets from the focal plane (bottom left, top right). DECODE and AttentionUNet provide the visually most appealing reconstructions. In the high-density regions (bottom left), the AttentionUNet reconstruction yields slightly less overlap of the approaching structures. Table 5 quantifies this observation. The AttentionUNet yields a better *RMSE* and *JI* than DECODE.

**Table 5.**
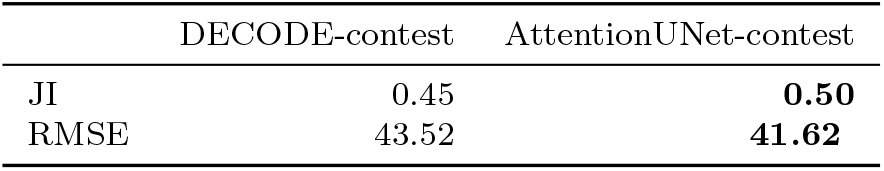
The finetuned AttentionUNet achieves the best Jaccard index and RMSE on the SuperResolution fightclub contest dataset for high emitter densities.

**Fig. 7.**
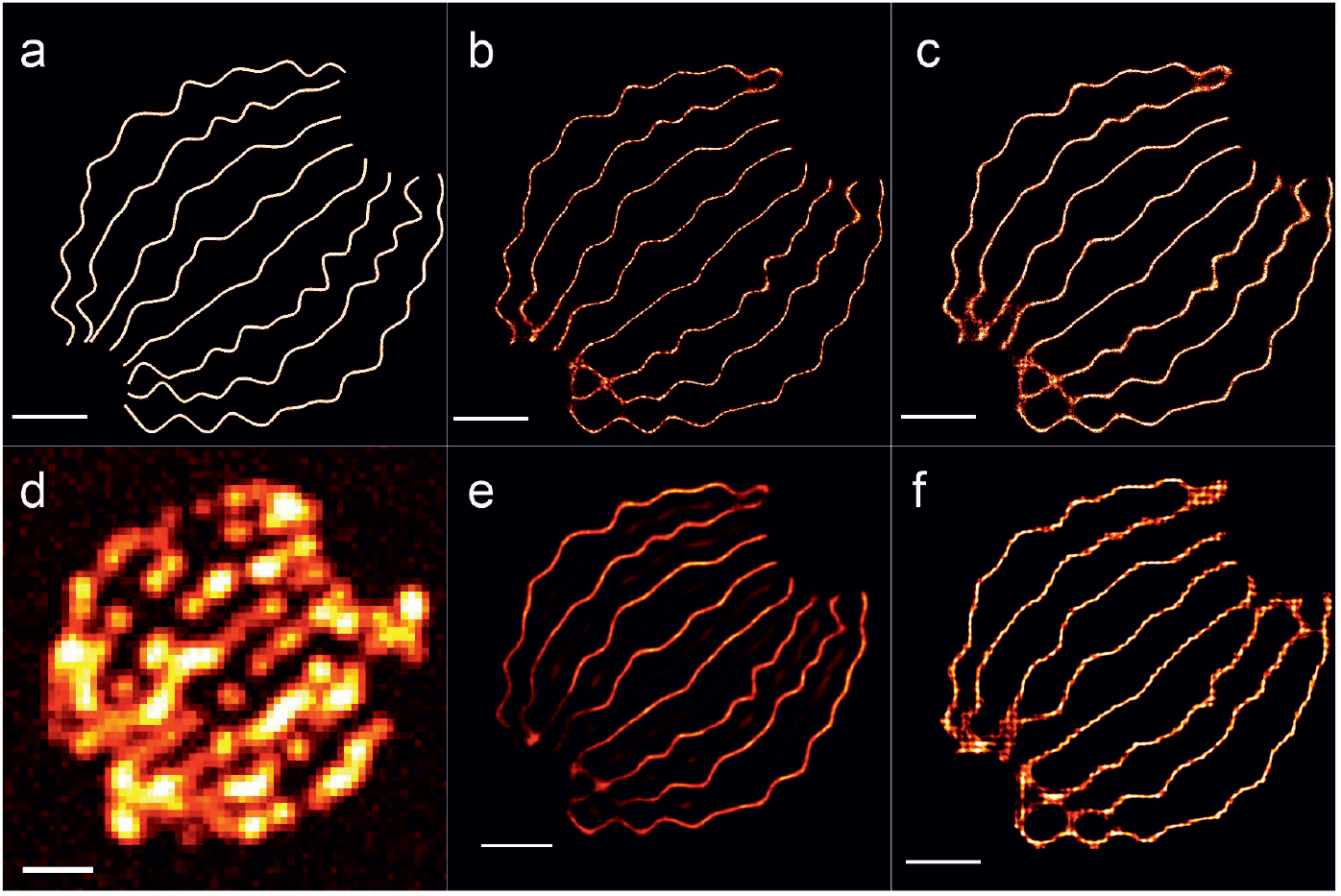
Contest. **a** Ground truth. **b** AttentionUNet(fine tuned). **c** DECODE (fine tuned). **d** Raw frame **e** SRRF. **f** Compressed Sensing (sig=180nm; lambda=250; it=1000). Scale bar = 1 µm

**Fig. 8.**
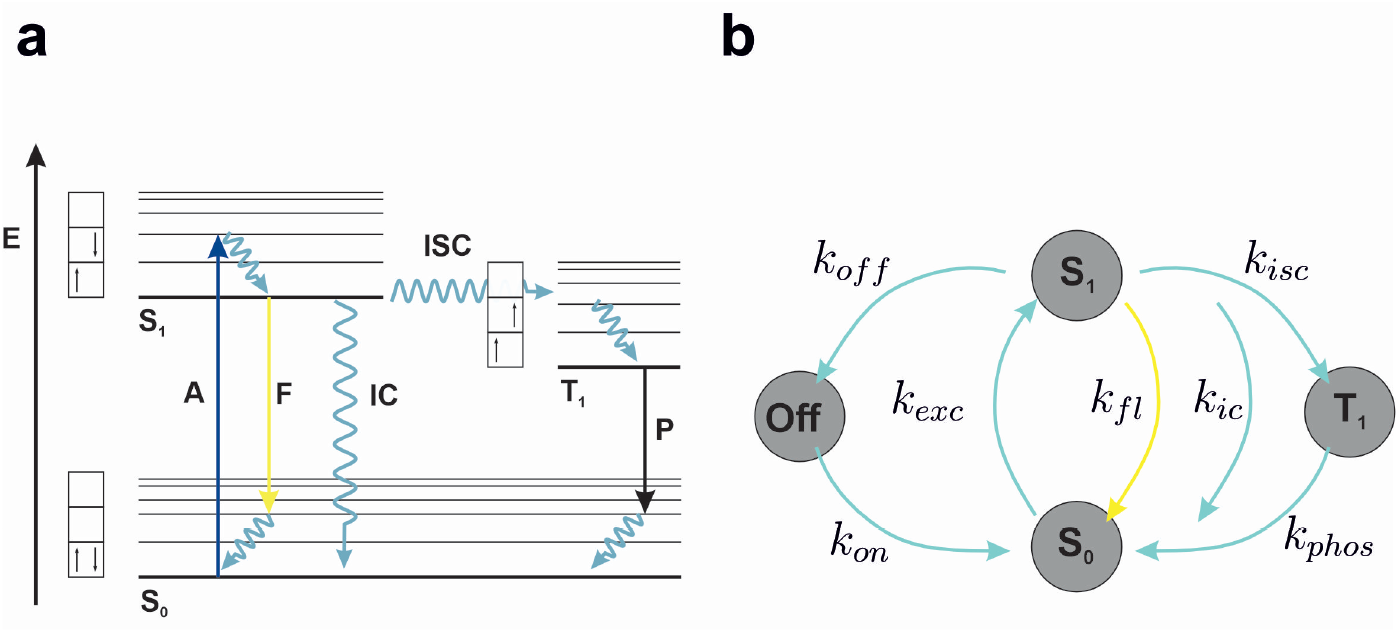
Transition model. **a)** Jablonski diagram of the photophysical states **b)** Resulting transition model.

**Fig. 9.**
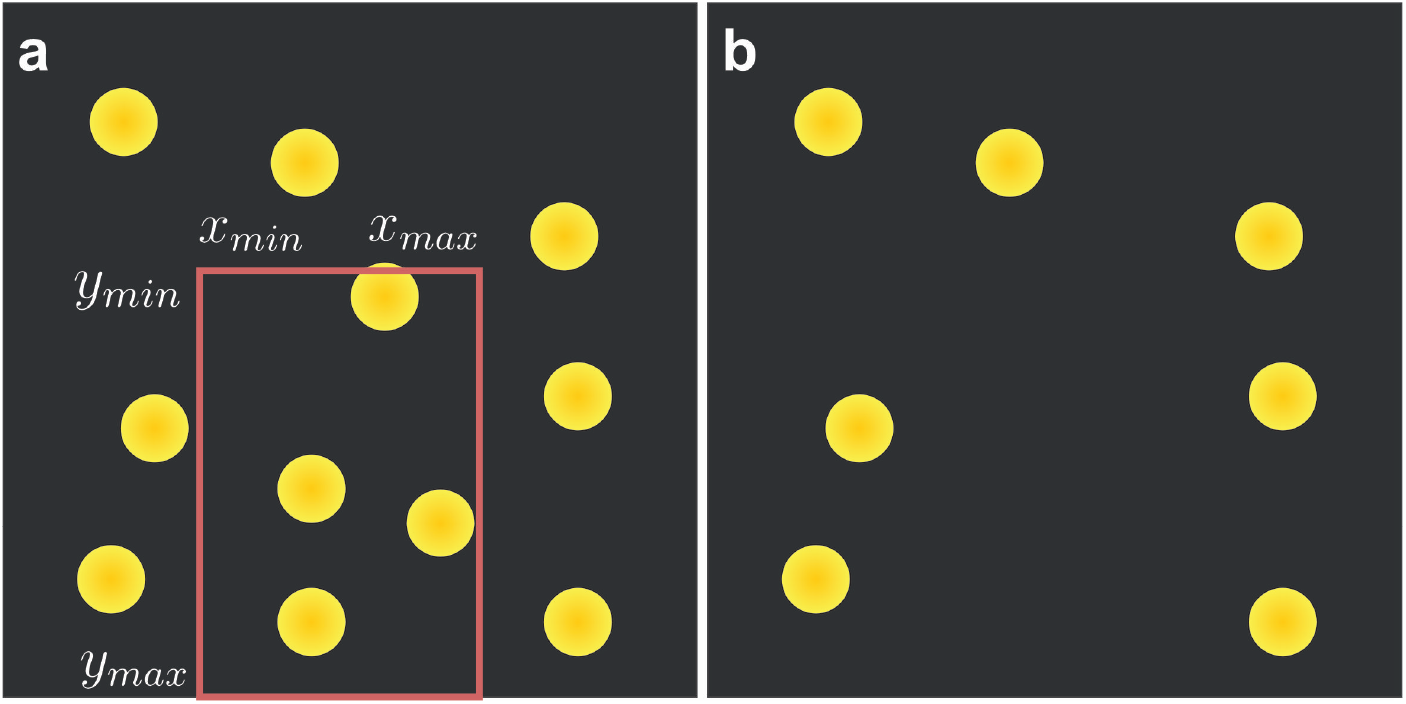
Dropout Box. **a)** Image before spatial dropout is applied. Coordinates for the dropout box are drawn from a uniform distribution 𝒰, i.e. *x*_*min*_ = min(***x***), where ***x*** = {*x*_1_, *x*_2_} ∼ 𝒰 (0, 1) multiplied with the respective image dimensions. **b)** Image after spatial dropout

**Fig. 10.**
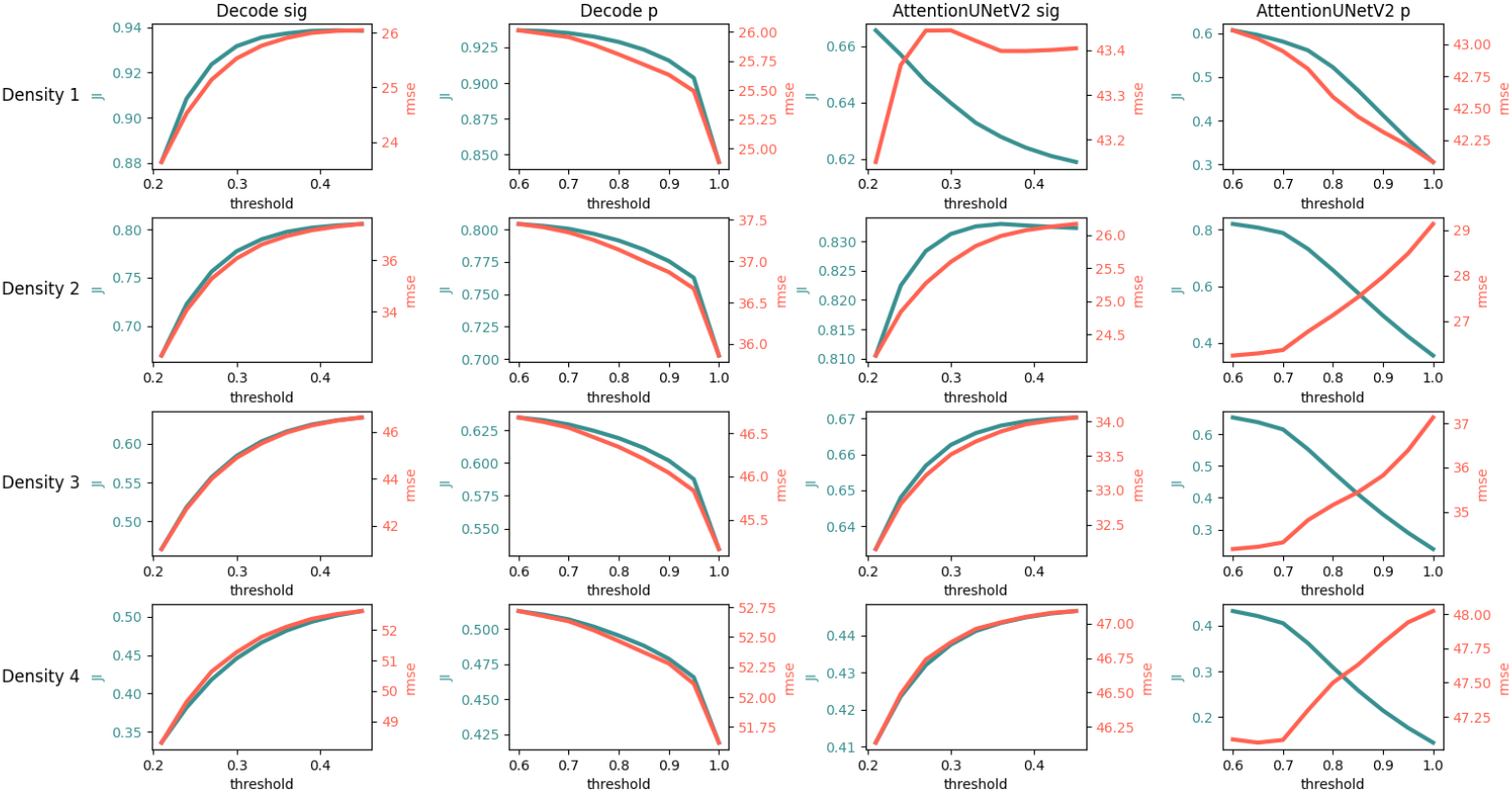
JI and RMSE under different sigma and p thresholds. Density is increasing from top to bottom, columns denote the different models.

Since these data contain no temporal context, we would expect a similar performance to DECODE. However, *JI* index and *RMSE* are slightly better. This leads to the hypothesis that neighboring emitters provide additional information about the overall structure in the case of filamentous structures, leading to improved positional accuracy.

## 3 Discussion

In this work, we use temporal correlations to gain additional information about the number and position of emitters in single-molecule localization microscopy. We present a deep neural network architecture that combines the previously used U-Net with multi-head attention to aggregate spatially encoded feature maps over a larger time context. To generate training data with time correlations, we implemented a Markov chain to simulate the behaviour of photo-physical states over long periods of time. These photon traces together with accurate noise models are used to generate training datasets with an extended temporal context of up to 50 frames.

A direct comparison of our method with current state-of-the-art fitters shows a significant improvement for emitters that are active over larger periods of time (Table 2). As shown in Figure 4, 76.59% of the fits are closer to the ground truth than the CRLB of the corresponding blinking event, compared to 39.95% for DECODE with a context size of three frames, and 17.53% for ThunderSTORM. We additionally found that the variation of this distance for the same emitter over the 50 frames timeframe is much smaller (1.85 nm) than for DECODE (4.35 nm) and ThunderSTORM (10.66 nm). While the difference between ThunderSTORM and DECODE reflects the difference in overall RMSE, the variation for our method is much lower than expected. This indicates that the network updates the emitter position with similar events over the whole timeframe, leading to similar positions and accuracy. The assumption that the improvement results from making use of the extended time context is further reinforced by a strong positive correlation between attention weights and *ON* events (Fig. 11).

**Fig. 11.**
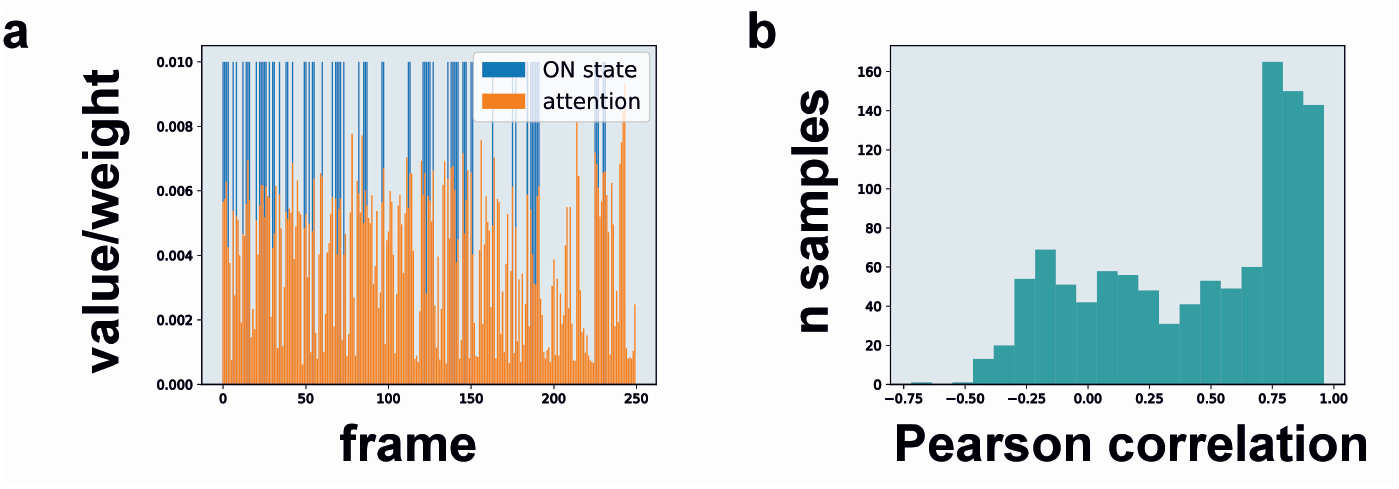
Temporal correlation of ON events and attention weights. **a** Time course of ON events (blue) and attention weights (orange). **b** Pearson correlation coefficient between ON state and attention weight.

Our approach works best under high-density conditions if temporal correlation is present in the data (Fig. 5, Sec 5). In this case, the multi-head attention update probably also finds nearby emitters that were previously active. For well-defined structures such as microtubules, this leads to an additional improvement of structural resolution.

As expected, increasing or decreasing the emitter density relative to the training data leads to reduced performance. If this discrepancy becomes too high, fine-tuning of the network is required. A possible explanation for this behavior might be that the implemented count loss constrains the network to the emitter density seen in the training data. Despite substantial efforts, we did not succeed in training a network that generalizes well for a large variety of different conditions. However, the current training (on different hidden dimensions) indicates that, similar to LLMs, better generalization could be achieved by upscaling the network, at the expense of increased requirements in terms of training data and computational resources.

The convergence of the training depends on the initialization of the network weights. Changing the random seed or the training data can lead to local minima that yield non-optimal reconstruction results. The loss function plays an important role for convergence, as it structures the optimization space. Possible improvements on this side could include normalization of the background loss, modifying the count loss to improve density generalization, and optimizing against distribution losses based e.g. on Kullback-Leibler divergence [31]. First experiments in this direction showed improved convergence at the cost of vastly higher computational requirements. Additional information might be gained from the photon count in the raw data, which is currently discarded by normalizing the network input.

The time for fitting using an already trained network is in the same range as for DECODE, and much faster than non-trainable algorithms (Table 6). Details on the time required for data simulation and training can be found in Materials and Methods. Further improvement of the performance could be achieved by increasing the temporal context up to the total length of an experiment, but the amount of parameters scales with the number of input frames as 𝒪 (*n*^2^), leading to a significant increase in computational requirements.

**Table 6.** Approximate inference time of the compared algorithms on the “Approaching lines” dataset.

|  | DECODE | AttentionUNet | ThunderSTORM (ME fitting) | SRRF | CS |
| --- | --- | --- | --- | --- | --- |
| ≈ time [s] | 31 | 35 | 3400 | 1000 | 10000 |

**Table 7.** Dataset statistics

| Name | frames<br>[count] | context<br>[frames] | density<br>[$\frac{emitter}{\mu m^2}$] | photons<br>[count] | ON-time<br>[frames] | structural<br>fill [%] |
| --- | --- | --- | --- | --- | --- | --- |
| Base | 100000 | 50 | 2.03 | 1764.47 | 1378.00 | 100 |
| Base test | 5000 | 50 | 2.03 | 1764.97 | 73.88 | 100 |
| Density 1 | 5000 | 50 | 0.38 | 1766.24 | 45.63 | 30 |
| Density 2 | 5000 | 50 | 0.73 | 1763.94 | 44.38 | 30 |
| Density 3 | 5000 | 50 | 1.12 | 1765.74 | 45.26 | 30 |
| Density 4 | 5000 | 50 | 1.52 | 1765.06 | 46.33 | 30 |
| CRLB 3 | 2500 | 50 | 0.18 | 1766.28 | 28.91 | 100 |
| CRLB 4 | 2500 | 50 | 0.17 | 1766.34 | 28.43 | 100 |
| Approaching<br>lines | 10000 | 50 | 1.50 | 1764.43 | 83.19 | 2 |
| Contest<br>(fine-tune) | 10000 | 50 | 4.72 | 1274.53 | 185.23 | 100 |
| ContestHD | 365 | — | $\approx 6$ | — | — | — |

On the “Approaching lines” dataset, our method yields a similar JI but improved RMSE compared to DECODE (Section 4). SRRF and CS show artifacts as the lines approach each other. It has to be noted that CS, while being able to distinguish emitters at high density, performs poorly under difficult noise conditions, especially when the noise level varies across the image. The SRRF algorithm depends on the radial symmetry of emitters and their intensity fluctuation over time. While highly fluctuating noise levels already pose a problem, it is even more difficult to distinguish closely spaced emitters on a line. The width of the line is reduced, and upon getting close together, both lines build a gradient maximum in the middle, merging the reconstruction into a single line with peak intensity at the center.

We tested both networks on the popular ContestHD dataset of the super-resolution fight club [8]. Since this dataset does not include temporal context, we expected our network to perform similarly to DECODE, but observed an improved JI and RMSE. This could either be due to our algorithm being more robust against changes of the PSF, or due to the structural context over the larger timeframe, which updates emitters towards other localisations within the structure as discussed above for the CRLB experiments, thus improving the precision of individual emitters “by accident”. In comparison, the SRRF reconstruction again displays mirroring artifacts between the lines that likely originate from a gradient maximum between both lines. The CS reconstruction suffers from pixelation artifacts and low precision for PSFs that are located further away from the focal plane, which can be explained by the change of the corresponding sigma.

As discussed above, the extended time context is used by the multi-head attention mechanism in our network to access not only intensity correlations, but also structural correlations that are distributed over a larger time window in the case of filaments. Structural correlations are also learned by deep neural networks to improve sparse localization maps [32, 33]. In a recent approach [34], time-context recurrent neural networks were applied to series of independently reconstructed localization images to infer super-resolved dynamics. Combining this method with our time-context aware fitting could be a promising approach, as it leverages correlations both from intensity fluctuations as well as from the spatial domain.

Current limitations of our approach include pixelation artifacts for increased uncertainty under very high noise and density conditions, and the restriction to twodimensional localization data. Although 3D fitting has not been implemented yet, it could easily be added using an according PSF model. The main limitation of current deep learning fitting methods including our approach is the requirement for training or fine-tuning for new data with slightly different conditions. Here, non-trainable methods such as SRRF have a clear advantage. We include a fast simulation engine with precise photophysics and accurate noise modeling together with instructions how to fine-tune or train the network from scratch. In the future, the training process could be made easier by inferring the input parameters for the simulation directly from the data, or completely avoided in a closed-loop approach using generative AI that learns to reproduce the experimental data from the encoded feature maps.

## 4 Conclusion

We present a new deep learning based framework for fitting high-density SMLM data using temporal context. The ground truth for training and fine-tuning is generated by a fast simulation engine with accurate noise models and photophysics. By using multi-head attention, our network is able to identify relevant frames and use the corresponding information to update emitter positions. Evaluations in comparison to the CRLB show that the precision of the fit scales with the amount of accessible information, i.e. with the number of frames in the context. The network outperforms other approaches with smaller time context for high-density and high-noise datasets if the conditions are similar to the training dataset. If conditions deviate too much from training conditions, fine-tuning is necessary. We also observed improved performance on data with no temporal context, which could be explained by structural correlations present over multiple frames. We hope that our work, for which we provide all source code and trained models, provides new insight into the nontrivial temporal context of SMLM data, and can serve as basis for further work.

## 5 Methods

### 5.1 Simulation engine

#### Background images

are created by creating an array (60×60) of random integers in the range ∈ [0, 255]. This array is convolved with a large sigma (5,5) Gaussian kernel. We create 10^4^ of those and randomly select one for each image.

#### Random localisations

are created as Nx2 array of randomly generated floats. The floats in the range ∈ [0.0, 1.0] are multiplied with the image size. We sample 3 × 10^7^ localisations as a pool to select from.

#### Structured localisations

are created by taking random points from within a binary image. The coordinates are normalized to the desired image dimension. Thus the input image should have a very high resolution to prevent discretized positions.

#### Subset selection

Each batch of 250 images contains a subset of the simulated localisations. The amount of emitters is computed with the desired approximated density and the average off time.

Points are augmented with a random box dropout, i.e., we create a random window where all emitters are discarded to prevent the network from learning densities.

#### Photoswitching

behavior for the drawn emitters is computed with a Markov chain model with rate matrix:

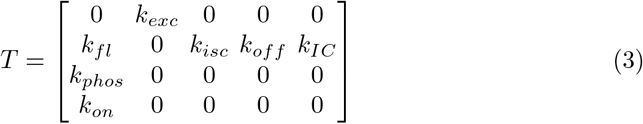

where *k*_*exc*_ is the excitation rate, *k*_*fl*_ the fluorescence rate, *k*_*IC*_ the internal conversion rate, *k*_*isc*_ the inter-system crossing rate and *k*_*off*_ the transition rate to the off-state. *k*_*isc*2_ and *k*_*on*_ denote the rate of return to the fluorescent ground state.

#### Transitions

are drawn from a categorical distribution:

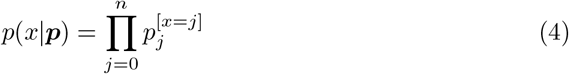

where *x* = *j* evaluates to 1 if *x* equals *j* and 0 if not. *p*_*j*_ is computed by:

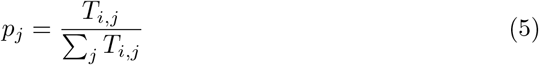

The lifetime of the state in row *i* is sampled from an exponential distribution:

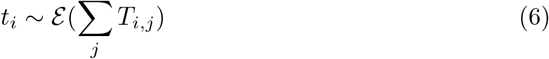

As a direct consequence of these transitions, the photon emission process of a fluorophore follows a sub-Poissonian statistic (single photon source antibunching).

#### Photon shot noise

Emitted photons are described by a wave function. The probability distribution of this wavefunction is the PSF. The area integral over the pixel 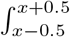*PSF* is approximated by the probability value at the center of the pixel. The probability value is multiplied by the expected number of photons. Summing the PSFs of all currently active emitters and adding a background signal yields the expected values for all pixels in the image. The number of photons per detector pixel can be described with a Poisson statistic. The probability for each photon to yield an electron is given by the quantum efficiency.

One could think there is a difference in whether the wavefunction collapses before an electron is excited (joint distribution of binominal and Bernoulli distribution) or after (expected value multiplied by the quantum efficiency *E*_*Q*_ before sampling Poisson). However, wave-particle duality states that both processes should be identical. This can be proven by computing the marginal distribution (law of total probability) of electrons *p*(*x*) collected given *n* input photons:

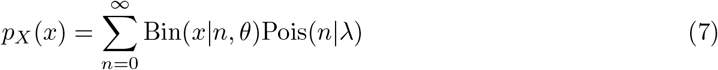

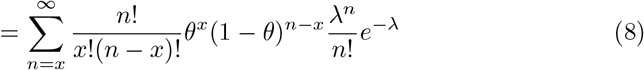

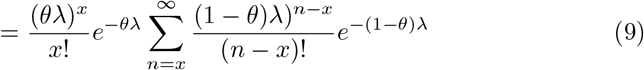

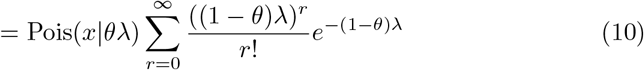

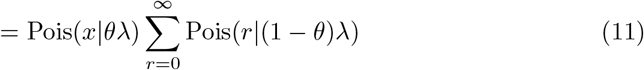

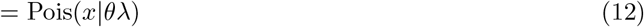

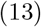

where *θ* = *E*_*Q*_.

#### Dark noise

arises due to thermal fluctuations within the sensor’s semiconductor material and its surrounding electronics. Electrons have a certain probability of spontaneously being excited into the conduction band. This leads to a certain number of events that occur even when no light is present. Dark noise can be quantified by the number of events *D*_*s*_ during the exposure time *t*. The process can be described as:

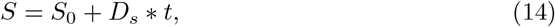

where *S*_0_ describes the true number of events, and *S* the output signal.

#### Amplification noise

occurs due to a probability smaller than one for an electron to trigger another electron in the avalanche photodiode. This can be described by a Gamma distribution

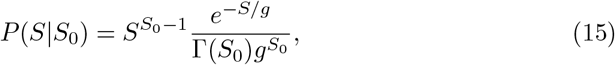

where Γ(*S*_*i*_) = (*S*_*i*_ − 1)! denotes the Gamma function and *g* denotes the gain factor.

#### Readout noise

occurs during the conversion of electrons into an electronic signal and is normally distributed. This normal distribution *N* of the electric current *I* with mean *I*_0_ and standard deviation *σ*_read_ is

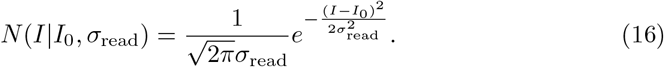

### 5.2 Metrics

#### GMM loss

For our loss function, we used a Gaussian Mixture Model as described in DECODE [10]. Here, each pixel of the feature map is able to predict one localisation. The first feature map yields the probability *p* to include a localisation at the center of the pixel with spatial shift ***µ*** = (*x, y*)^*T*^ deduced by feature map two and three. Together with the uncertainty values from feature maps four and five, we create a multivariate Gaussian distribution for each pixel:

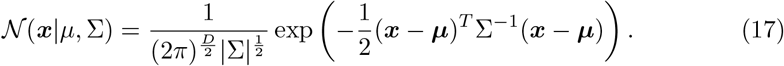

where *D* denotes the dimensionality and Σ the covariance matrix. Minimizing the negative log-likelihood of the multivariate Gaussian weighted by the probability *p*_*n*_,

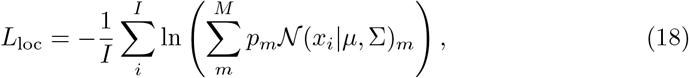

of this model for a given ground truth point *x*_*i*_, pushes *µ* towards the ground truth while minimizing the covariance matrix Σ:

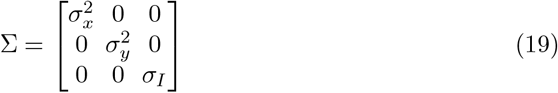

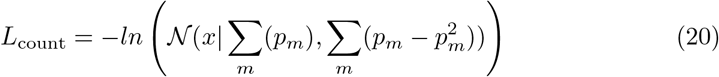

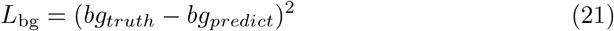

#### Jaccard Index

The Jaccard Index is a standard metric defining the percentage of true positive *T*_*P*_ localisations divided by the sum of true positives, false negatives *F*_*N*_ and false positives *F*_*P*_ :

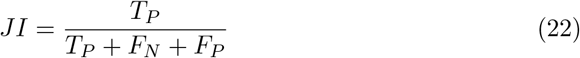

A localisation is counted as true positive if its distance is within 100 nm of a ground truth emitter. If multiple localisations are close to a ground truth emitter, the closest one is counted as *T*_*P*_ while the others are counted as *F*_*P*_.

#### RMSE

The Root Mean Squared Error (RMSE) qunatifies the distance of emitters to the ground truth and is defined as:

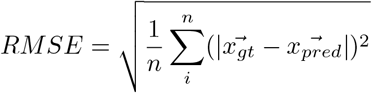

#### Efficiency score

The efficiency score introduced by [8] combines the RMSE with the JI into a single metric:

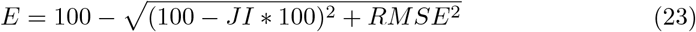

### 5.3 Network architecture

Artificial neural networks for sequential data such as time series differ in their architecture from the traditional fully connected multi-layer perceptron or convolutional neural networks. They include a hidden state vector to model memory of past elements of the sequence. Recurrent neural networks (RNN) suffer from the problem that the range of the memory is limited. LSTMs include additional “forget” gates to selectively remember different elements, but suffer from the same problem. A more general concept is attention, which can learn relationships between far-away elements of a sequence.

The attention mechanism in its current form was first described in the paper “Attention is all you need” [20], where it was shown to achieve outstanding results for sequence processing. The mechanism is composed of a linear mapping to the components Query (Q), Key (K) and Value (V) that takes an embedding of position-encoded tokens such as words or image patches and learns to map each element to other positions in the sequence. One popular application is the prediction of the next word in a text, which is the basis of large language models such as GPT.

The process can be described by the following mathematical equation:

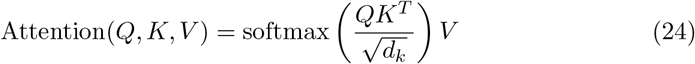

where *d*_*k*_ describes the embedded dimension size and 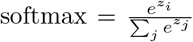 for the i-th row in the matrix.

Figure 11 a shows the relationship between the ON state of an emitter and the attention weights of the underlying pixel for the first frame. It can be seen that the network pays more attention to frames with signal. We surmise that the network is able to collect additional positional information over these frames. To support that assumption, we computed the Pearson correlation of ON state and attention weights for all emitters in a batch of our test dataset.

### 5.4 Diffusion Network

**Fig. 12.**
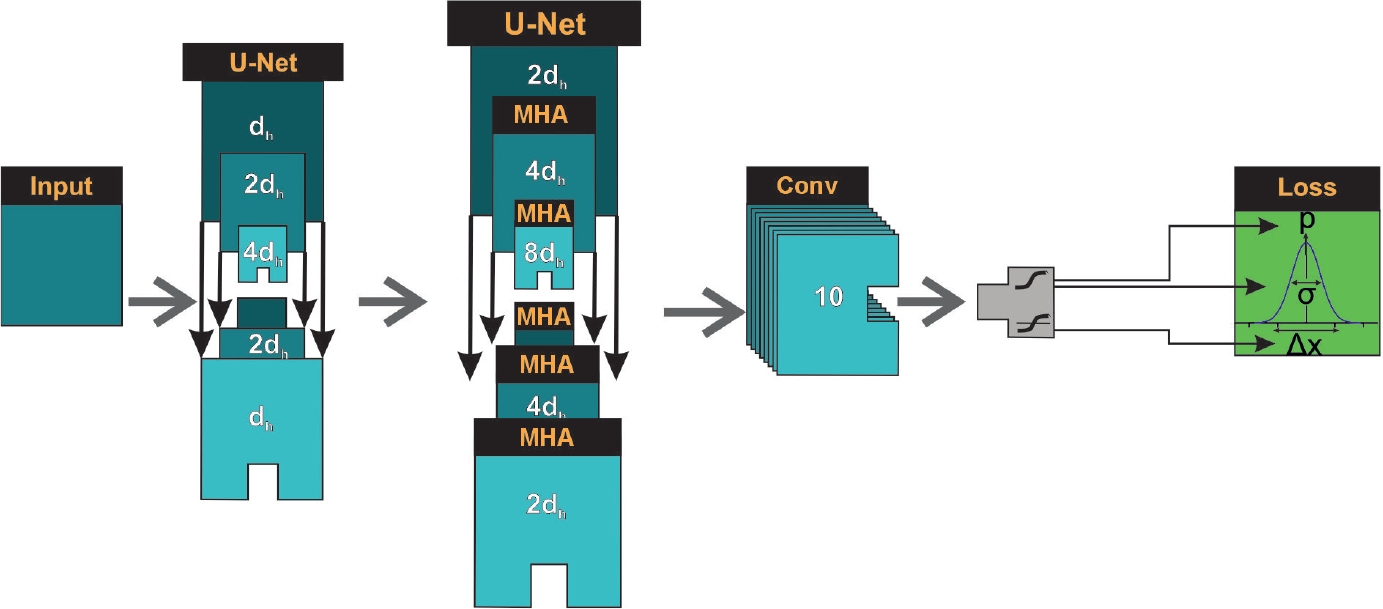
Diffusion Network. Network architecture with multi-head attention (MHA) in the second U-Net.

### 5.5 Training

We trained the network with 500 batches of 50 images of 60×60 px for 20 epochs. The training was performed on an Nvidia GeForce 4090 GTX. Total training time from scratch was ≈24 hours. The batch size is limited by the hardware. As shown in Figure 3, the lightest network is composed of a simple U-Net but also delivers the worst performance. Adding a consecutive attention block to the U-Net mainly influences the background loss, i.e., the network is better in distinguishing signal from noise. An improvement in positional loss is only achieved after adding a second consecutive U-Net.

#### 5.5.1 Base training

Since the base training requires extensive resources and multiple steps of optimization it can, in contrast to the fine-tuning, not be identically reproduced. Training with the given parameters and datasets should however yield similar results. We provide a checkpoint to our base training under the link at the end of the methods section.

#### 5.5.2 CRLB training

The datasets “CRLB 3” and “CRLB 4” can be reproduced with the provided .yaml files. We split the trainingsset “CRLB 4” into a training and validation set with a ratio of 80-20. Note that batches of 50 frames stay together and unshuffled to preserve switching behavior. We compute an efficiency score every two iterations. To fine-tune the base training, we optimized the parameters with optuna for 20 trials, maximizing the efficiency score. This leads to the parameters in the training .yaml files. Using these files reproduces the used training checkpoints.

### 5.6 Training data

The .yaml configuration files in the GitHub repository provide full reproducibility of the datasets. Running simulate images.py sets all random seeds to a specific value, resulting in exactly the same dataset as used in the paper.

### 5.7 Availability of code and data

Code to create training data, train networks and predict localization positions as well as a trained base model are available under https://github.com/super-resolution/SMLM-AttentionAI

